# Developmental accumulation of gene body and transposon non-CpG methylation in the zebrafish brain

**DOI:** 10.1101/2020.12.17.423365

**Authors:** Samuel E Ross, Daniel Hesselson, Ozren Bogdanovic

## Abstract

DNA methylation predominantly occurs at CG dinucleotides in vertebrate genomes; however, non-CG methylation (mCH) is also detectable in vertebrate tissues, most notably in the nervous system. In mammals it is well established that mCH is targeted to CAC trinucleotides by DNMT3A during nervous system development where it is enriched in gene bodies and associated with transcriptional repression. However, the conservation of developmental mCH accumulation and its deposition by DNMT3A is largely unexplored and has yet to be functionally demonstrated in other vertebrates. In this study, by analyzing DNA methylomes and transcriptomes of zebrafish brains, we identified enrichment of mCH at CAC trinucleotides (mCAC) at defined transposon motifs as well as in developmentally downregulated genes associated with developmental and neural functions. We further generated and analyzed DNA methylomes and transcriptomes of developing zebrafish larvae and demonstrated that, like in mammals, mCH accumulates during post-embryonic brain development. Finally, by employing CRISPR/Cas9 technology, we unraveled a conserved role for Dnmt3a enzymes in developmental mCAC deposition. Overall, this work demonstrates the evolutionary conservation of developmental mCH dynamics and highlights the potential of zebrafish as a model to study mCH regulation and function during normal and perturbed development.

## Background

In genomes of vertebrate adult somatic cells, the majority of CpG sites are methylated (>80%) with the exception of CpG-rich promoters and distal regulatory elements (Bird, 2002; Jones, 2012; Schübeler, 2015). While otherwise ubiquitous, CpG methylation (mCG) at regulatory elements is known to participate in long-term gene silencing processes (de Mendoza et al., 2019). In mammals, albeit at much lower levels, methylation of cytosines outside the CpG context (mCH, H = T,C,A) has also been reported in the majority of tissues (Schultz et al., 2015). mCH, or more particularly methylation of CA dinucleotides (mCA), occurs most commonly in mammalian embryonic stem cells (ESCs) and in the brain (Lister et al., 2009, 2013; Schultz et al., 2015). In ESCs, mCH is enriched at CAG trinucleotides in gene bodies and is positively correlated with gene expression. Additionally, increased levels of mCH were observed at repetitive elements in ESCs (Ziller et al., 2011; Arand et al., 2012; Guo et al., 2014b). In mammalian brains, mCH rivals the levels of mCG and is enriched at CAC trinucleotides (mCAC) in gene bodies where it negatively correlates with expression and is deposited de novo by DNMT3A (Lister et al., 2013). In line with its repressive role in the nervous system, mCH is depleted at open chromatin regions (Lister et al., 2013). mCH in the postnatal mammalian brain displays a rapid increase during initial phases of synaptogenesis, which corresponds to two to four weeks in mouse, and first two years of life in humans. This is followed by a longer period of slower accumulation (Lister et al., 2013).

While mCH is found at high levels and studied extensively in plants (Zhang et al., 2018), the function of mCH in vertebrates is less well known. Several studies, however, have demonstrated that Methyl-CpG Binding Protein 2 (MeCP2) is able to bind to and regulate genes marked by mCH, which was particularly evident at long genes (Guo et al., 2014a; Chen et al., 2015; Gabel et al., 2015; Boxer et al., 2020; Clemens et al., 2020). Whether this is due to biological or technical reasons is currently debated (Raman et al., 2018). Mutations in MeCP2 are the most prevalent cause of Rett syndrome, and interestingly, altered readout of mCH deposited by DNMT3A appears to play a central role in Rett syndrome pathogenesis (Chen et al., 2015; Lavery et al., 2020). MeCP2 is conserved across vertebrates, such as zebrafish, where depletion of MeCP2 results in similar pathologies to Rett syndrome including altered motor behavior, improper synapse formation and acute inflammation (Pietri et al., 2013; Gao et al., 2015; Nozawa et al., 2017; van der Vaart et al., 2017).

A recent report described the conserved enrichment of mCH in vertebrate brains, which originated alongside MeCP2 and DNMT3A enzymes at the root of the vertebrate lineage (de Mendoza et al., 2020). This study also highlighted the anti-correlation between gene body mCH and expression in some, but not all, vertebrate brains. In our previous work, we found highly specific mCH enrichment at TGCT tetranucleotides within zebrafish mosaic satellite repeats in embryonic and adult tissues, deposited by the teleost specific Dnmt3ba enzyme (Ross et al., 2020). However, the developmental dynamics and distribution of neural-specific mCH, and a functional role for DNMT3A or MeCP2 in relation to mCH, has yet to be demonstrated outside of mammalian brains. Here we expand upon the utility of the zebrafish model in the study of mCH and reveal that like in mammals, mCH accumulates during brain development via Dnmt3a enzymes and becomes enriched at downregulated genes and Tc1-like transposable elements. This study thus extends our knowledge of vertebrate mCH conservation and lays the foundation for future work that will allow for the precise dissection of mCH regulatory functions during embryogenesis and nervous system formation.

## Materials and Methods

### Zebrafish usage and ethics

Zebrafish work was conducted at the Garvan Institute of Medical Research in accordance with the Animal Ethics Committee AEC approval 20/09 and with the Australian Code of Practice for Care and Use of Animals for Scientific Purposes. Adult wild type (AB/Tübingen) *Danio rerio* (zebrafish) were bred in an equal ratio of males and females. Embryos were collected 0 hours post-fertilization (hpf) and incubated in 1X E3 medium (5 mM NaCl, 0.33 mM CaCl2, 0.17 mM KCl, 0.33mM H14MgO11S) for four days at 28.5°C before being transferred onto a filtered system.

### Genomic DNA and RNA extraction

Whole brains were dissected from zebrafish larvae and adults before being snap-frozen in liquid nitrogen and stored at −80°C. Genomic DNA (gDNA) was extracted from brains using the QIAGEN DNeasy Blood & Tissue Kit (QIAGEN, Chadstone, VIC, Australia) according to manufacturer’s instructions. For RNA extraction, half of the lysate from the first step of DNA extraction from the QIAGEN DNeasy Blood & Tissue Kit was added to TRIsure (Bioline) and purified following manufacturer’s instructions. All experiments in this study were performed in two biological replicates.

### CRISPR/Cas9 zebrafish knockouts

Guide RNAs (gRNA) targeting *dnmt3aa* and *dnmt3ab* loci were designed with CRISPRscan (Moreno-Mateos et al., 2015). gRNAs for both loci were synthesized and co-injected into 1-cell stage embryos as previously described (Ross et al., 2020). CRISPR/Cas9 knockouts (cKO) fish were grown to four weeks of age before their brains were harvested for DNA and RNA extraction. Amplicons surrounding the CRISPR/CaS9 cut sites were PCR-amplified from genomic DNA, ligated to NEXTFLEX Bisulfite-Seq barcodes (PerkinElmer, Waltham, MA, USA), and spiked into libraries that were sequenced on the Illumina HiSeqX platform. Knockout efficiencies were calculated from the sequenced amplicons using CRISPResso (Pinello et al., 2016). RNA was reverse transcribed to cDNA using the SensiFAST™ cDNA Synthesis Kit (Bioline), following the manufacturer’s protocol and subjected to qPCR analysis. Relative expression levels were calculated using the 2−ΔΔCT method and *bactin* gene as the control transcript. Two sample t-tests were performed on CT values. All oligos used in this study can be found in **Supplemental table 1**.

### Whole genome bisulfite sequencing (WGBS)

WGBS libraries were prepared from 500 ng of zebrafish brain gDNA and spiked with 0.025 ng of unmethylated lambda phage DNA (Promega, Madison, WI, USA) as previously described (Ross et al., 2020).

### Reduced representation bisulfite sequencing (RRBS)

RRBS libraries were prepared from 500 ng of zebrafish brain gDNA spiked with 0.025 ng of unmethylated lambda phage DNA (Promega, Madison, WI, USA). gDNA was digested with 10U BccI and 10U SspI for 2h. A separate aliquot was digested with 20 U MspI (New England BioLabs, Ipswich, MA, USA). RRBS libraries were constructed as previously described (Ross et al., 2020), sequenced, and the BAM files corresponding to different digestion reactions (BccI/SspI or MspI) were merged before downstream analysis.

### WGBS and RRBS data analyses

WGBS reads were trimmed with Trimmomatic: ILLUMINACLIP:TruSeq3-PE.fa:2:30:10 SLIDINGWINDOW:5:20 LEADING:3 TRAILING:3 MINLEN:60 (Bolger et al., 2014), and mapped using WALT (-m 5 -t 20 -N 10000000) (Chen et al., 2016) onto the GRCz11 reference genome (UCSC), containing the λ genome. BAM files were deduplicated using sambamba markdup (Tarasov et al., 2015). RRBS data were processed as above with the additional option of: HEADCROP:5 CROP:140 added during trimming and without deduplication. BAM files were made FLAG-compatible and processed with CGmapTools (Chen et al., 2016; Guo et al., 2018) (convert bam2cgmap) to obtain ATCGmap files, which were corrected for CH positions that showed evidence of CG SNPs (de Mendoza et al., 2020). Genomic data were visualized in UCSC (Kent et al., 2002) and IGV (Robinson et al., 2011) browsers.

### DNA sequence motif analyses

BED file coordinates of the 10,000 most highly methylated mCH and mCAC sites, with a minimal depth of 10, were extended by 4 base pairs upstream and downstream. The resulting files were used as input for HOMER “findMotifsGenome.pl” function (Heinz et al., 2010), establishing the search for de novo motifs of length 9 (-len 9 -size given) with the GRCz11 genome used as the background sequence. Motifs were visualized using the “ggseqlogo” package in R (Wagih, 2017) and motif positions in the genome were called using the scanMotifGenomeWide.pl function (with and without - mask option checked).

### mCH level calculation and plotting

Bedgraphs were generated from corrected CGmapTools outputs and converted to bigWig using bedGraphToBigwig script from Kent utils. Average mCH levels were determined from bedGraph files and calculated by dividing the sum of reads supporting a methylated cytosine by the sum of all reads mapping to that position. mCH levels in genomic features and gene bodies were calculated using BEDtools map (Quinlan and Hall, 2010). mCH levels, TPMs, and gene length were plotted using the boxplot function in R (outline = FALSE). Heatmaps were generated using deepTools (Ramirez et al., 2014) computeMatrix with the following parameters: “computeMatrix scale-regions -m 650 -b 500 -a 500 -bs 25”. NAN values were replaced with 0. Heatmaps were plotted with the plotHeatmap function and sorted and clustered based on methylation levels. Scatterplots were generated using the geom_bin2d function in ggplot2 ((bins=50) + geom_smooth(method=lm)). Pearson correlations were calculated using the *rcorr* function.

### Repeatmasker track analyses

Repeatmsker tracks were obtained from UCSC. The percentage of repeat subfamilies overlapping the top-methylated CAC motifs was determined with BEDtools (intersectBed).

### ChIP-seq Analysis

Brain H3K27ac fastq files (Kaaij et al., 2016) were trimmed with Trimmomatic (ILLUMINACLIP:TruSeq3-SE.fa:2:30:10 SLIDINGWINDOW:5:20 LEADING:3 TRAILING:3 MINLEN:20) (Bolger et al., 2014) before they were mapped to the GRCz11 genome using bowtie2 with default settings (Langmead and Salzberg, 2012). BAM files were deduplicated using sambamba markdup (Tarasov et al., 2015). H3K27ac peaks were called using MACS2 (Zhang et al., 2008).

### RNA-seq

RNA-seq libraries were prepared with 1000 ng of input RNA using the KAPA mRNA HyperPrep Kit, according to manufacturer’s instructions.

### RNA-seq analyses

RNA-seq reads were trimmed using Trimmomatic: ILLUMINACLIP: TruSeq3-PE.fa:2:30:10 SLIDING-WINDOW:5:20 LEADING:3 TRAILING:3 MINLEN:60 (Bolger et al., 2014) and aligned to the GRCz11 genome using STAR (Dobin et al., 2013). Differential gene expression analysis was performed using edgeR (Robinson et al., 2010). (Robinson et al., 2010) with genes selected based on a minimum ±1.5 logFC (FDR <0.05) between any of the analyzed time points. Z-scores were calculated based on the log2 transformations of TPM values and plotted with the *pheatmap* package in R. Analysis of published RNA-seq data was performed based on the provided read count tables with TPM values calculated from the average of five, six-month old brain datasets (Aramillo Irizar et al., 2018) or from collated counts of 30 neurons (Lange et al., 2020).

## Results and discussion

### mCH is enriched at defined CAC-containing motifs in zebrafish brains

To investigate mCH in the zebrafish nervous system, we analyzed WGBS data (bisulfite conversion >99.5%) of adult brain, as well as of adult liver, to use as a non-neural control tissue (Bogdanovic et al., 2016; Skvortsova et al., 2019). We employed stringent genotype correction (de Mendoza et al., 2020) to allow for more sensitive interrogation of mCH patterns. To better understand the sequence context of mCH deposition in the zebrafish brain and how it compares to non-neural tissues (liver), we performed a motif search on the 10,000 most highly methylated CH sites. Expectedly, we recovered the TGCT motif associated with mosaic satellite repeats in both brain and liver (Ross et al., 2020), and the previously characterized TACAC-containing motif (Lister et al., 2013; de Mendoza et al., 2020), which was specific to the brain sample (Supplemental Figure 1A). Furthermore, the brain sample showed a notable enrichment in CAC trinucleotide methylation (~1.5%) with a 3-fold increase compared to the unmethylated lambda genome spike-in control (~0.5%) (Supplemental Figure 1B). This mCAC enrichment was not evident in the liver sample, thus eliminating the possibility of sequence-specific biases or artifacts pertaining to bisulfite conversion (Olova et al., 2018) (Supplemental Figure 1B). We next investigated the genomic distribution of mCAC in zebrafish brains to assess if the depletion in regulatory elements and enrichment in gene bodies previously described in mammals (Lister et al., 2013; Guo et al., 2014a; He and Ecker, 2015) is evolutionarily conserved. To achieve this, we annotated transcription start sites (TSS), exons, introns, 5’UTRs, 3’UTRs, intergenic regions, and sites of H3K27ac enrichment, which correspond to active gene-regulatory elements (Kaaij et al., 2016). We found that intronic and intergenic regions are the only regions enriched in mCAC and mCH (Supplemental Figure 1C-E), whereas a notable depletion, similar to the one described in mammals, was observed at H3K27ac peaks (Supplemental Figure 1C-F). These results support the previous observations of mCA presence in gene bodies of vertebrate brains (de Mendoza et al., 2020) and demonstrate a conserved depletion of mCH in other genomic features, such as in active regulatory regions.

Further analyses of sequence motifs associated with the mCAC context unraveled two novel sequences in addition to the previously described vertebrate-conserved TACAC motif (Figure 1A). Due to previous associations of mCH with repetitive DNA in zebrafish (Ross et al., 2020), we wanted to assess if any of the most significantly methylated CAC motifs were enriched in repetitive elements. The top TACAC motif displayed comparable methylation levels in repetitive elements and in the repeat-masked genomic fraction, with an average methylation level nearly 3-fold higher (~4%) than the average global mCAC levels (Figure 1B). However, for the remaining two motifs we found a robust increase in average methylation levels at repetitive elements when compared to the repeat-masked genome (~6.5% and 6%, Figure 1B). Further analysis revealed that the TACAC motif is broadly distributed in the genome whereas the second and third motif were mainly located in TDR and TC1DR3 repetitive elements, respectively (Figure 1C). We also found that average methylation of the top three methylated CAC motifs correlated strongly with overall mCH in gene bodies (*R*=0.52) (Figure 1D), suggestive of a significant contribution of CAC methylation to gene body mCH. This correlation was stronger than the one observed between average gene mCH and gene length (*R*=0.36), which was previously described in mammals (Gabel et al., 2015; Boxer et al., 2020) (Supplemental Figure 1G). Additionally, when Gene ontology (GO) analysis (Raudvere et al., 2019) of genes containing methylated TDR and TC1DR3 CAC motifs was performed we found overrepresentation in developmental and neural development terms (Figure 1E). Due to its widespread genomic abundance the TACAC motif was omitted from the GO analysis. Overall, zebrafish brain is enriched in mCH, particularly in the CAC trinucleotide context, predominantly in introns and intergenic regions, as well as in members of the Tc1-mariner transposon family.

**Figure 1.**
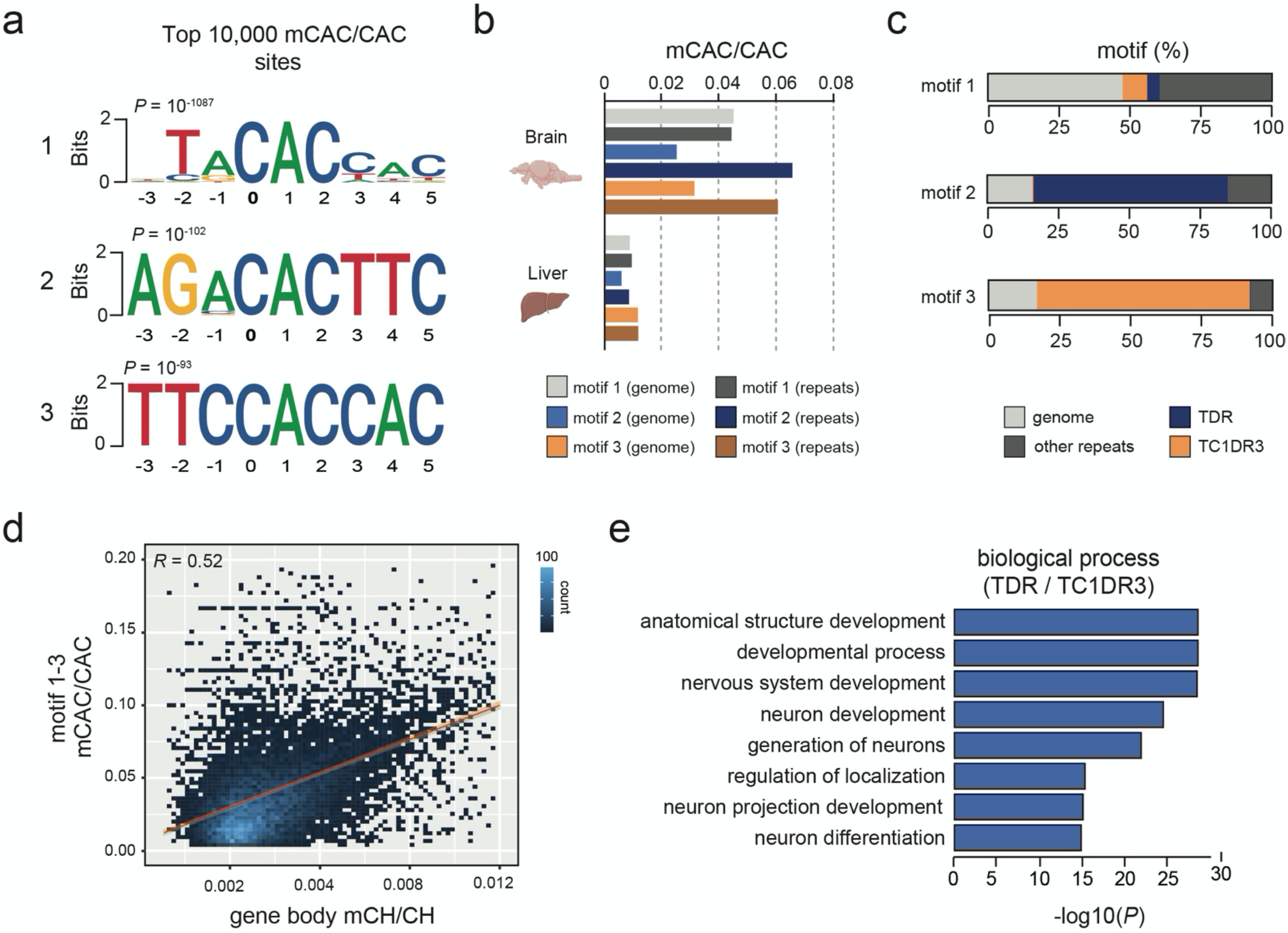
mCH is enriched at defined CAC-containing motifs in zebrafish brains. **a)** Top three motifs called from the 10,000 most methylated CAC trinucleotides in the zebrafish brain. **b)** Average mCAC/CAC methylation levels of the top three mCAC motifs in the bulk genome and repetitive elements of zebrafish brain and liver. **c)** Genomic annotation of top three mCAC motifs. **d)** Genomic correlation between average gene body mCH/CH and mCAC/CAC at top three most methylated CAC motifs. **e)** Gene ontology enrichment of genes containing methylated CAC motifs in TDR and TC1DR3 elements.

### mCH is targeted to genes downregulated in the nervous system

To explore the relationship between mCH and gene expression in zebrafish, we plotted average gene mCH and mCAC values against gene expression (transcripts per million-TPM) levels from available datasets (Bogdanovic et al., 2016; Aramillo Irizar et al., 2018) (Figure 2A). We revealed a gene cluster (n=1,860) with higher-than-average mCH levels, which displayed lower expression than the bulk transcriptome (Figure 2A). To provide more genomic context to these findings, we first interrogated whether this elevated mCH was driven by a higher proportion of intron sequence in these genes. To that end, we plotted gene length, gene body mCAC, intron mCAC and exon mCAC for mCH-enriched genes as well as for genes that did not display any notable mCH enrichment (Figure 2A, B). While the mCH-enriched genes were significantly longer, in line with observations in mammals (Gabel et al., 2015; Boxer et al., 2020), the elevation in mCH was not driven exclusively by intron contribution as both introns and exons located within these genes had significantly higher levels of mCAC (Figure 2B). Interestingly, genes with higher levels of mCAC also contained a higher percentage of Tc1-like elements in relation to total gene length (Figure 2C). To confirm the observation of poorly expressed genes being marked by mCH, we plotted average TPM levels from total RNA-seq data from adult brains (Aramillo Irizar et al., 2018) and combined single cell data from neurons (Lange et al., 2020) (Figure 2D). Genes with higher levels of mCH (cluster 1) had significantly lower average TPM when compared to genes with moderate/low levels of mCH (cluster 2), or to a randomly selected subset of genes (n=1,860, Figure 2D). This difference in expression levels between the two clusters was even more pronounced in neurons where mCH is expected to be the highest based on mammalian data (Lister et al., 2013) (Figure 2D). GO analysis of these highly mCH-methylated genes again revealed enrichment for terms associated with embryonic and neural development, and notably, 53% of these genes contained methylated TDR/TC1DR3 motifs compared to only 13% of genes with moderate/low mCH levels (Figure 2E, Supplemental Figure 2A). This enrichment of developmental genes for mCH is in line with observations in vertebrate brains (de Mendoza et al., 2020). Therefore, like in mammals, mCH in the zebrafish adult brain is associated with transcriptional repression which is particularly evident in long genes.

**Figure 2.**
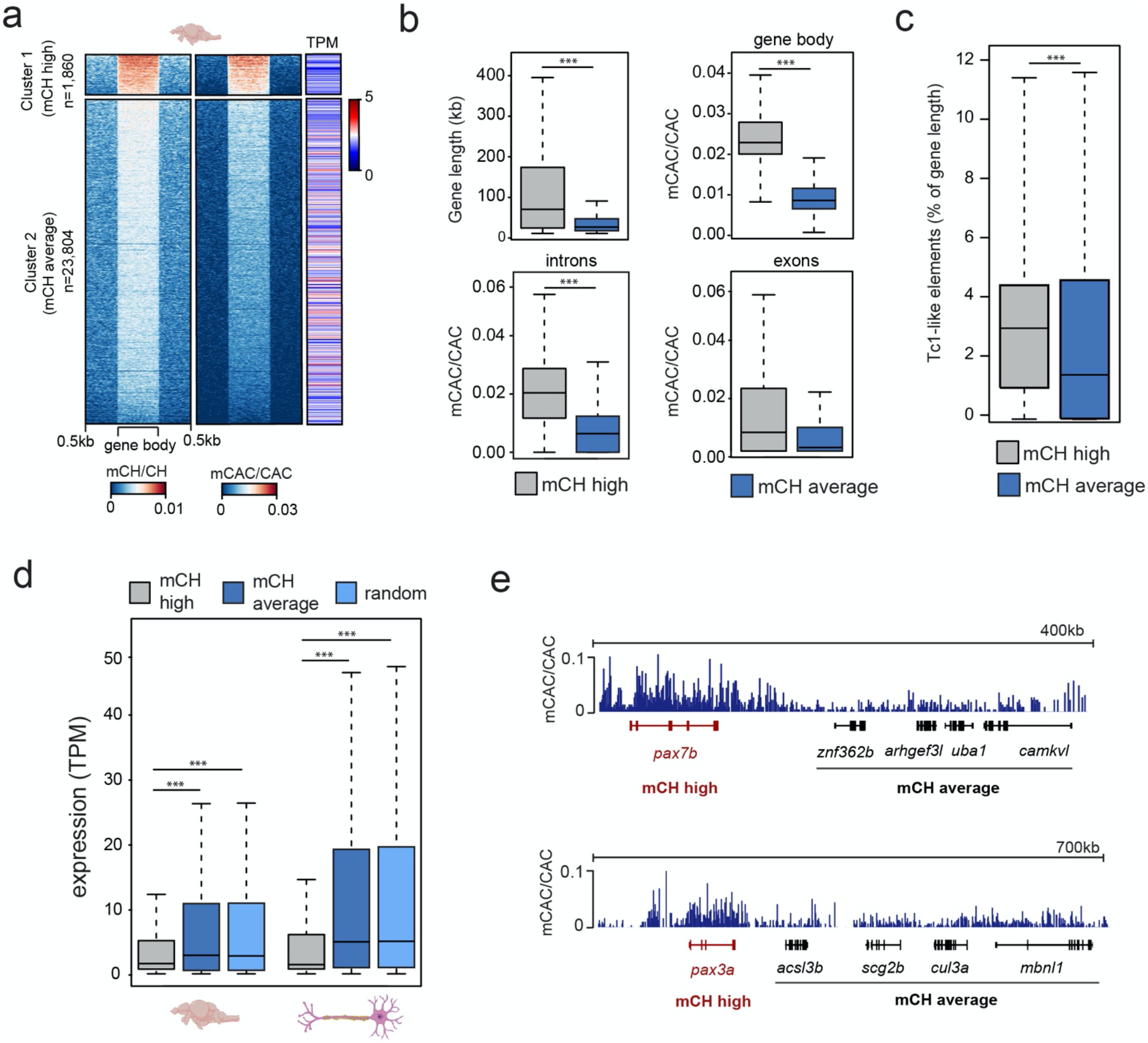
mCH is present at long genes with low expression levels. **a)** mCH/CH levels, mCAC/CAC levels, and gene expression represented as transcripts per million (TPM) in adult zebrafish brains. **b)** Comparisons of gene length (top left), average gene mCAC/CAC (top right), average exon mCAC/CAC (bottom left) and average intron mCAC/CAC (bottom right) in genes with high (cluster 1) and moderate/low levels of mCAC/CAC (cluster 2) (Wilcoxon test, *** *P* <0.001). **c)** Tc1-like element percentage of total gene length in genes with high (cluster 1) and moderate/low levels of mCAC/CAC (cluster 2) (Wilcoxon test, *** *P* <0.001). **d)** Average expression levels (TPM) in six-month old brains (n = 5) and neurons (n = 30) at genes with high (cluster 1), or moderate/low (cluster 2) mCH, and a random subset of genes (n=1,860) sampled from cluster 2 (Wilcoxon test, *** *P* <0.001). **e)** Genome browser snapshot of mCAC/CAC levels in cluster 1 (red) and cluster 2 (black) genes in adult brains.

### non-CG methylation accumulates during zebrafish brain development

As mCH has previously been shown to accumulate during mammalian brain development (Lister et al., 2013), we next investigated whether comparable mCH dynamics could be observed in zebrafish. We generated RRBS libraries (Meissner et al., 2005) using a combination of enzymes to enrich for regions containing highly methylated CAC motifs (Figure 1A) identified in adult brains. We assayed zebrafish brains starting from one week, where brain structures such as the cerebrum are not identifiable, up until six weeks and adulthood, where all structures are discernable (Maeyama and Nakayasu, 2000). This analysis revealed a gradual increase in mCAC in the brains of one-week to six-week-old zebrafish followed by a more notable increase in adult brains (Figure 3A). RNA-seq analysis across this period also revealed a gradual decrease in the expression of components of DNA methylation machinery as cells presumably become more differentiated (Figure 3B). Moreover, *mecp2* expression increased during brain development coinciding with increase in mCH. This observation supports a conserved role for MeCP2 in the regulation of genes marked by mCH in vertebrates. Differential expression analysis of all genes across brain development revealed two major gene clusters which were either consistently upregulated (cluster 1) or downregulated (cluster 2) during brain development (Figure 3C). Interestingly, 18% of the downregulated genes belonged to the mCH-enriched gene cluster, compared to only 4% of the upregulated genes. Downregulated genes were also associated with terms related to cell division and development while genes that were upregulated were enriched in terms associated with adaptive immunity (Figure 3D). These results are consistent with ongoing developmental processes in the larval brain and with the notion that the adaptive immune system of zebrafish does not develop until four to six weeks post fertilization (Lam et al., 2004).

**Figure 3.**
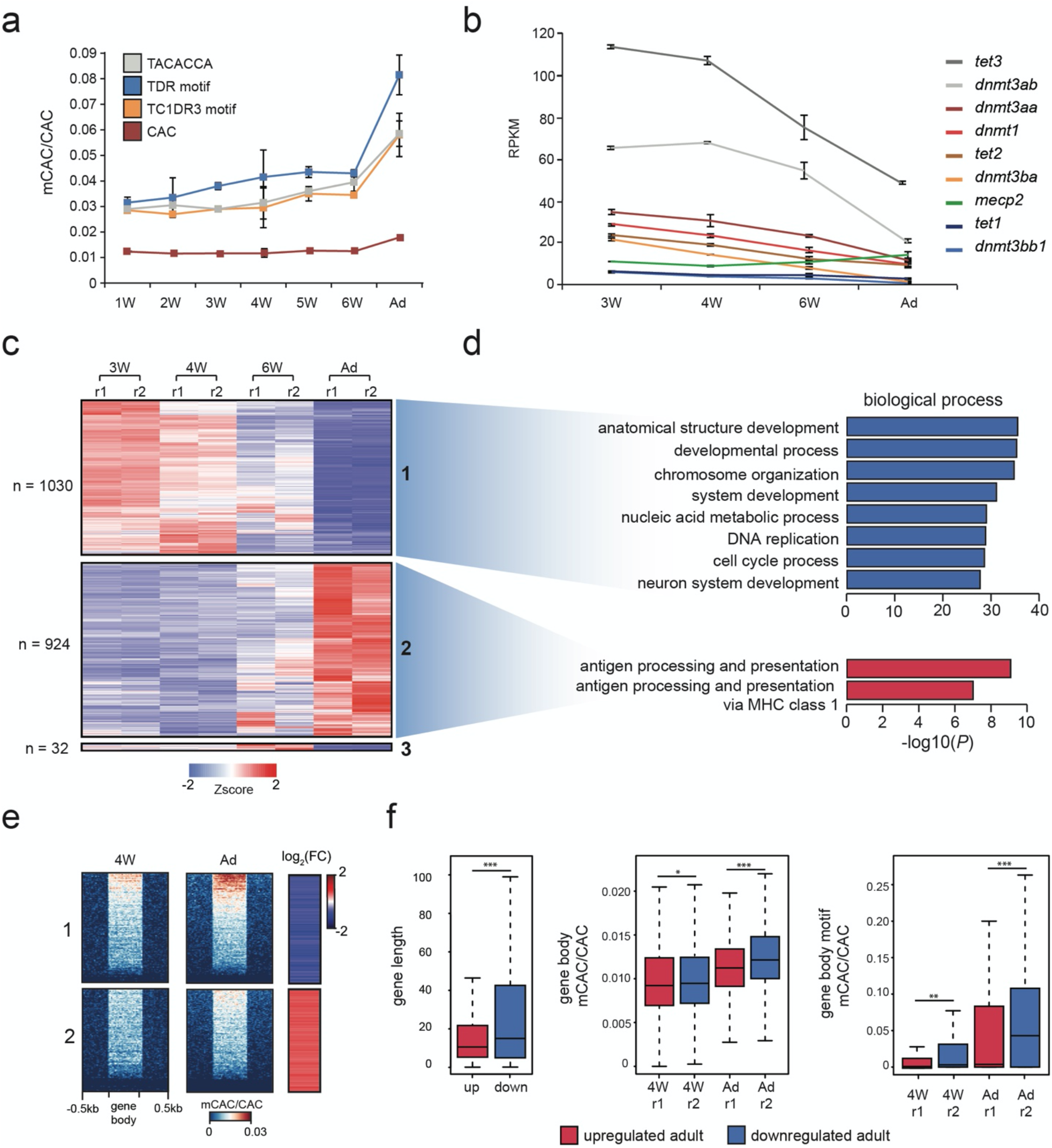
mCH accumulates in the developing nervous system. **a)** mCAC/CAC levels at all CAC trinucleotides and top methylated CAC motifs in larval (W = weeks old) and adult (Ad) brains, as determined by RRBS. Data is represented as the average of two biological replicates with error bars (standard deviation). **b)** RPKM (reads per kilobase per million) values of *dnmt*, *tet*, and *mecp2* transcripts in larval and adult brains determined by RNA-seq. Data is represented as the average of two biological replicates with error bars (standard deviation). **c)** Transcription intensities of a merged collection of differentially expressed genes called between all pairwise comparisons of larval and adult stages (r1 - replicate 1, r2 = replicate 2). **d)** Gene ontology enrichment of differentially expressed genes in larval and adult brains. **e)** mCAC levels and relative RNA expression levels (log2 fold change 4W / Ad) at differentially expressed genes. **f)** Comparisons of gene length (left), gene body mCAC/CAC (middle), and gene body mCAC/CAC at top methylated motifs (right) in genes that are either upregulated or downregulated in the adult brain (Wilcoxon test, * *P* <0.05, ** *P*<0.01, *** *P* <0.001).

To understand better how mCH and gene expression dynamics track over developmental time, we generated WGBS datasets for four-week-old brain tissue and compared these data against adult brain WGBS and RNA-seq data. Analysis of mCAC levels of differentially expressed genes revealed that developmentally downregulated genes accumulate more mCH when compared to upregulated ones. (Figure 3E). This trend was also observed when visualizing mCAC levels and gene expression levels across all genes (Supplemental Figure 2B, D). Furthermore, quantification of mCAC levels at all mCAC trinucleotides and at the highly methylated motifs (all three combined), confirmed a significant increase in the methylation of developmentally downregulated genes (Figure 3F). This increase in mCH in adult brains is uncoupled from global mCG changes, as global mCG decreases over developmental time during nervous system development (Supplemental Figure 2D,E) (Bogdanovic et al., 2016). Altogether, these results demonstrate robust anticorrelation between mCH and gene expression during brain development in zebrafish, as well as developmental mCH accumulation, similar to the observations in mammals.

### Dnmt3a enzymes are required for methylation of CAC trinucleotides in the zebrafish brain

Finally, to investigate if Dnmt3a-dependent methylation of CAC trinucleotides is evolutionarily conserved in zebrafish, we generated dnmt3aa/*dnmt3ab* CRISPR/Cas9 double knockouts (cKO). qPCR analysis of cDNA extracted from the brains of four-week-old cKOs revealed a 50% reduction in expression levels for both *dnmt3aa* and *dnmt3ab* (Figure 4A). Sequencing of amplicons surrounding the CRISPR/Cas9 cut sites demonstrated comparable estimates of genome editing efficiency (Figure 4B). Finally, WGBS analysis of these samples showed that depletion of *dnmt3aa*/*dnmt3ab* resulted in a significant (*P* < 0.05, Wilcoxon test) reduction in global mCAC levels as well as in a notable depletion (43%) in the methylation of top mCAC motifs (*P* < 0.01, Wilcoxon test) (Figure 4C). The reduction in mCAC levels in these cKOs can be observed across the majority of gene bodies (Figure 4D) and on a genome- and locus-specific scale (Figure 4E). Notably, these perturbations in mCH did not result in any obvious morphological defects in the cKO fish. Finer analysis of brain morphology, behavior, and potential inflammatory processes in these animals will be a focus of future studies. Additionally, no changes in global CpG methylation levels or significant DMRs could be detected (data not shown) between the cKO and WT brain, suggestive of mCH deposition being a major function of Dnmt3a enzymes in zebrafish. However, given the incomplete KO, the possible redundancies with other zebrafish DNMTs (Goll and Halpern, 2011), the propagation by Dnmt1 once established, and the use of bulk cell DNA methylome data, we cannot completely rule out a role for these enzymes in neuronal CpG methylation. Nevertheless, the clear role in mCH deposition, and the fact that Dnmt3a enzymes can be traced back to the root of vertebrates (de Mendoza et al., 2020), suggests conserved functional importance of mCH in the vertebrate nervous system.

**Figure 4.**
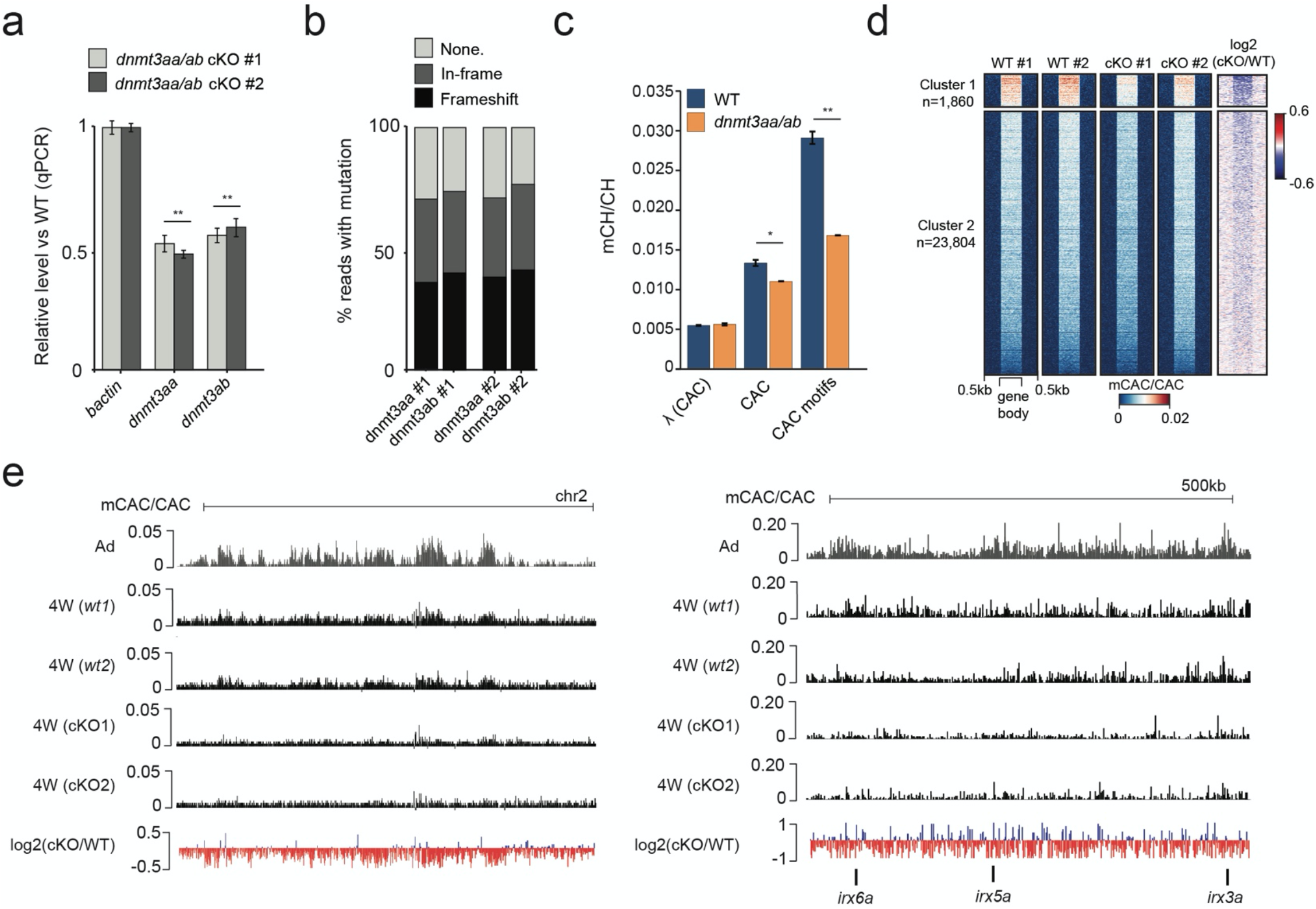
Dnmt3a enzymes are required for methylation of CAC trinucleotides in the zebrafish brain. **a)** Transcript levels of *dnmt3aa* and *dnmt3ab* in four-week-old *dnmt3aa/ab* CRISPR/Cas9 KO (cKO) zebrafish brains relative to wild type (WT). The data is represented as the mean of technical replicates with error bars showing the standard error (two sample *t*-test, ** *P* < 0.01). **b)** Percentage of reads with no mutation, in-frame mutations, or frameshift mutations, which map to *dnmt3aa* and *dnmt3ab* loci in *dnmt3aa/ab* cKOs. **c)** Average mCAC/CAC levels at all CAC trinucleotides and the top methylated CAC motifs in four-week-old WT and *dnmt3aa/ab* cKO brain. The data is represented as the average of two WGBS biological replicates (Wilcoxon test, * *P* <0.05, ** *P* < 0.01, λ = unmethylated lambda spike in control). **d)** mCAC/CAC levels of all gene bodies in four-week-old WT and *dnmt3aa/ab* cKO brains. **e)** Genome browser snapshot of mCAC/CAC levels in adult brains, four-week-old WT brains and four-week-old *dnmt3aa/ab* cKO brains.

## Discussion

While mCH has been established as an important base modification with likely biological functions during mammalian brain development (Lister et al., 2013), and links to Rett syndrome pathogenesis (Chen et al., 2015; Gabel et al., 2015; Boxer et al., 2020; Lavery et al., 2020), there are still many unknowns related to its regulation and function. Furthermore, the evolutionary conservation of mCH, the mCH “writer” - DNM3TA, and the mCH “reader” - MeCP2 in vertebrates suggests that these regulatory pathways could have an ancestral role in vertebrate neurobiology (de Mendoza et al., 202a).

In the current manuscript, we describe the conservation of developmental mCH dynamics in the zebrafish nervous system. In zebrafish, like in mammals, mCH is enriched at CAC trinucleotides in gene bodies where it accumulates during brain development. Also, in line with observations in mammals, mCH depletion is evident at H3K27ac-marked regulatory regions. Similarly to our recent work on TGCT methylation of mosaic satellite repeats in zebrafish (Ross et al., 2020), and previous reports of mCH enrichment at repetitive elements in mammals (Ziller et al., 2011; Arand et al., 2012; Guo et al., 2014b), we find high levels of mCH at defined motifs associated with Tc1-like transposons (~6%). This recurring observation of mCH enrichment at repeats in multiple species supports a possible role for mCH, or DNMT3A recruitment, in the regulation of repetitive elements. In mammalian brains, active transposition of repeat elements has been shown to drive mosaicism in neuronal genomes (Muotri et al., 2005; Macia et al., 2017; Bodea et al., 2018), while MeCP2 was described as a repressor of LINE-1 elements in mouse neurons (Yu et al., 2001; Muotri et al., 2010). These data thus suggest that mCH could play an important role in regulating repetitive elements in the vertebrate brain, particularly at CG-sparse regions or active repeats such as Tc1-like transposons. These observations are also reminiscent of mCH targeting by DNMT3-related enzymes in plants (Law and Jacobsen, 2010).

Finally, in the current study, we have generated transient CRISRPR/Cas9 KOs for *dnmt3aa*/*dnmt3ab* and demonstrated a conserved role for these enzymes in the deposition of mCH, and mCAC in particular, in the zebrafish nervous system. These KOs only had an obvious effect on mCH but not mCG levels, suggestive of a direct conservation for mCH in the vertebrate nervous system. Overall, this work provides novel insight into the evolutionary conservation of vertebrate mCH patterning and highlights the utility of the zebrafish model system, which is amenable to CRISPR/Cas9 screens, drug screens and developmental imaging, for the studies of mCH and brain development *in vivo*.

## Author contribution

OB conceived the study. SR performed bioinformatic analyses and CRISPR/Cas9 experiments. DH extracted brain samples. SR and OB wrote the manuscript. All authors contributed to, read, and approved the final manuscript.

## Acknowledgments

Images of zebrafish brain, liver, and neurons were created with Biorender. We thank Alex de Mendoza for critical reading of the manuscript.

## Availability of data and materials

Data generated for this submission have been uploaded to ArrayExpress https://www.ebi.ac.uk/arrayexpress/ under the accession number E-MTAB-9924.

---

**Supplemental figure 1.**
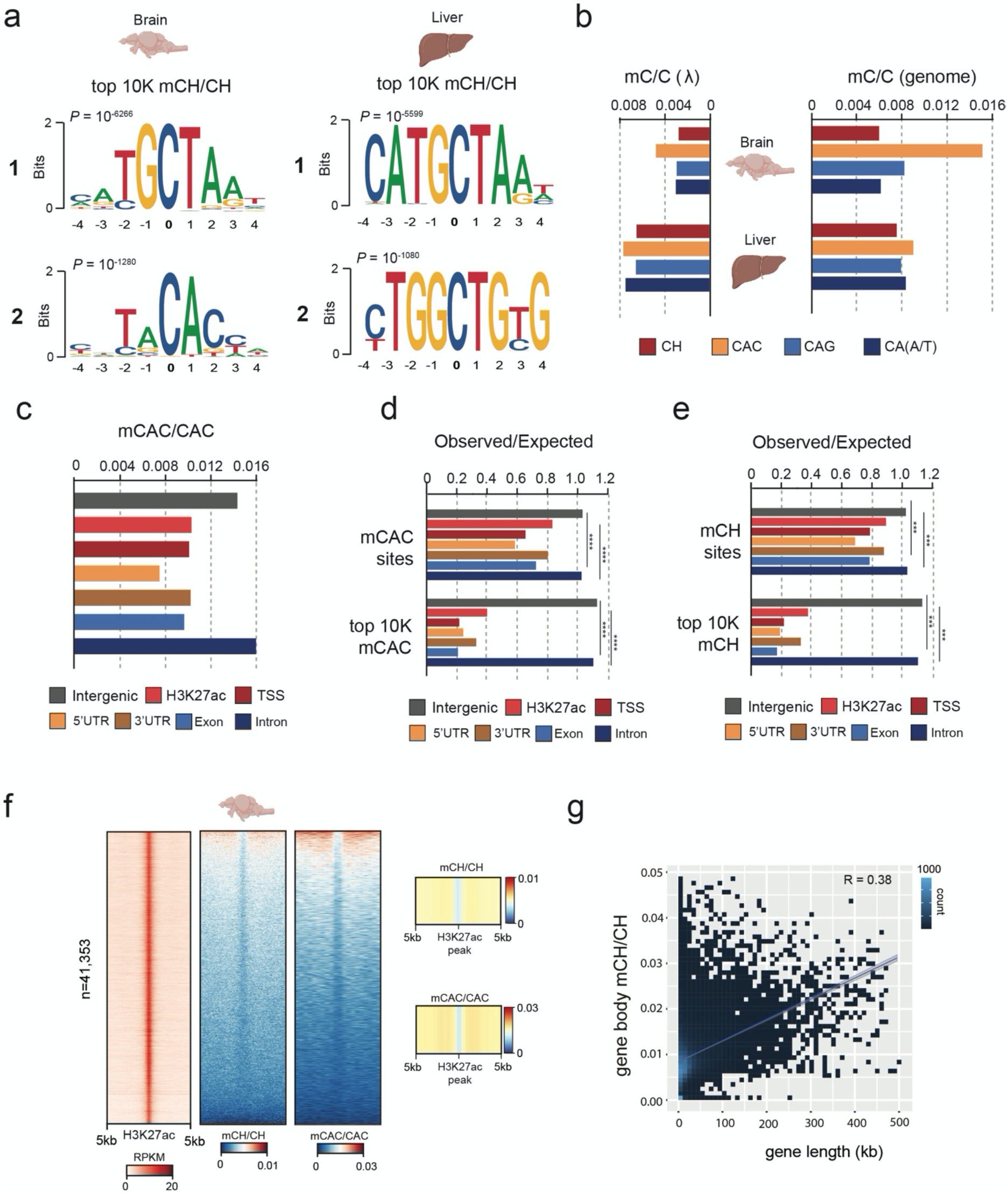
**a)** Top two motifs called from 10,000 most highly methylated CH dinucleotides in zebrafish brain and liver. **b)** Average non-CG methylation levels (mC/C) at CH, CAC, CAG and CA(A/T) nucleotides in brain, liver and unmethylated lambda spike-in controls (λ). **c)** Average CAC methylation levels across diverse genomic features in the zebrafish brain. **d)** Observed-over-expected ratios of all methylated CAC sites and the 10,000 most highly methylated CAC sites (χ² test *** *P* < 0.001). **e)** Observed-over-expected ratios of all methylated CH sites and the 10,000 most highly methylated CH sites (χ² test *** *P* < 0.001). **f)** H3K27ac enrichment expressed as reads per kilobase per million (RPKM), mCH/CH and mCAC/CAC in H3K27ac peaks in adult zebrafish brains. **g)** Genome-wide correlation between average gene body mCAC/CAC and gene length.

**Supplemental figure 2.**
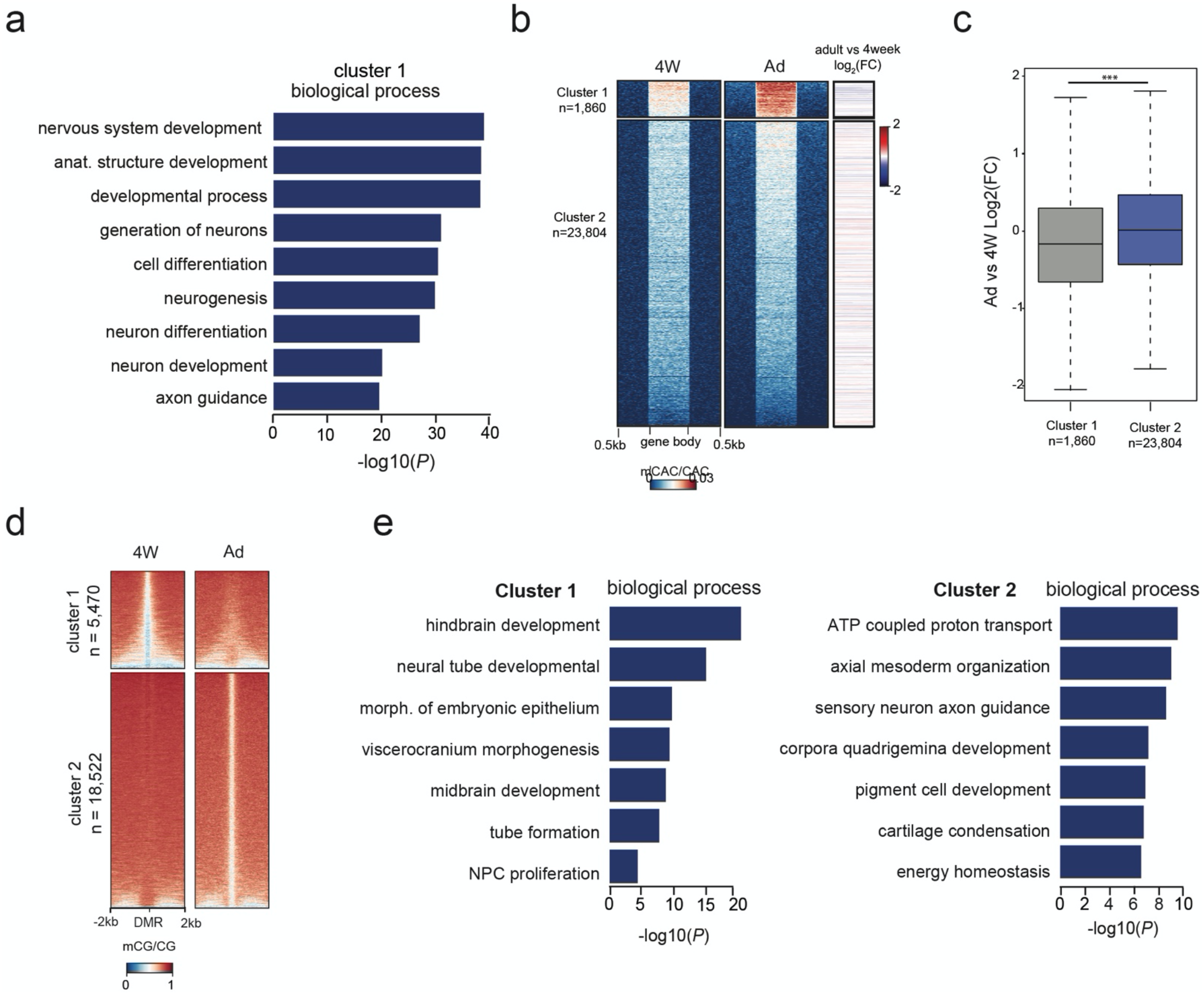
**a)** Gene ontology enrichment of genes with high levels of mCAC/CAC. **b)** mCAC levels and relative RNA expression levels (log2 fold change Ad/4w) at all genes. Positive fold change indicates upregulation in adult. **c)** RNA expression levels in adult and four-week-old brains (log2 fold change Ad/4w) in genes marked by high levels of mCH (cluster 1) vs other genes (cluster 2) (Wilcoxon test, *** *P* < 0.001). Positive fold change indicates upregulation in adult **d)** mCG/CG methylation levels at differentially methylated regions (DMRs) in four-week-old (4W) and Ad brain samples. **e)** Genomic region enrichment of hypomethylated regions in four-week-old brains (cluster 1) and hypomethylated regions in adult brains (cluster 2).

